# Analysis Of Respiratory Syncytial Virus Reveals Conserved Rna Secondary Structural Motifs And Impact Viral Lifecycle

**DOI:** 10.1101/2024.08.26.609688

**Authors:** Elena M Thornhill, Ryan J. Andrews, Zachary Lozier, Elizabeth Carino, Marie-Anne Rameix-Welti, Jean-François Eléouët, Walter N. Moss, David Verhoeven

## Abstract

An analysis that combined bioinformatics, comparative sequence/structural analysis, and experimental assays has been completed on respiratory syncytial virus (RSV). Both the genomic RNA and its reverse complement were studied using the novel bioinformatics pipeline ScanFold, which predicted 49 regions on RSV RNAs that appear to encode functional secondary structures (based on their unusually stable sequence order). Multiple motifs appear to be conserved between RSV and related virus strains, including one region within the F gene, which had a highly favorable overall prediction metric of a paired secondary structure. This motif was subjected to additional experimental analyses using SHAPE analysis to confirm ScanFold predicted secondary structure. In subsequent analysis, RSV F mRNA transcripts were made by in vitro transcription using T7 polymerase and transcripts which relaxed the predicted secondary structure yielded slightly higher mRNA transcripts and protein expression levels as wildtype F. However, using reverse genetics for comparison of viruses containing wildtype or relaxed F suggested that the predicted secondary structures may be critical for RSV replication in cells. To our knowledge, this is the first study to examine conserved RNA structures across multiple RSV strains and may help identify potential therapeutic targets to inhibit.

## Introduction

Respiratory Syncytial Virus (RSV) is a significant respiratory viral pathogen that circulates throughout human populations (64 million infections per year) with clinical symptoms manifesting very similarly to influenza virus. RSV typically causes up to 125,000 hospitalizations for bronchiolitis (1,2), 20.3% of the secondary pneumococcal pneumonia cases (3), and is the main viral catalyst for secondary middle ear bacterial infections. RSV is so ubiquitous that almost all children have been infected at least once by two years of age. RSV morbidity is not just restricted to the young. 10,000 deaths are associated with the virus in the elderly per year in the US alone (4). Despite some recent advances in immunogen design, there have been no efficacious vaccine for this virus (5) until recently with two vaccines (RSVpreF3 and RSVpreF from Glaxo Smith and Pfizer had been approved for the elderly (47,48). In a study of 72 Gavi-eligible countries RSV caused 1.2 million discounted DALYs, and US$611 million discounted direct costs (6) highlighting RSV’s impact on the world economy and population.

The RSV genome is a single-stranded non-segmented negative sense RNA ∼15,000 nucleotides in length. The genome encodes 10 genes and 11 different proteins. The 11th protein results from overlapping ORF’s in a mechanism that isn’t well understood. Transcription proceeds in order along the genome in a start stop mechanism, such that the NS genes should have the highest mRNA transcript levels and with the last gene in the genome L exhibiting the lowest levels of mRNA transcripts (49–50). Recently, a new study has shown a non-linear decline in gene transcript aboundance that is not uniform (51). Detection primer optimization for each RSV gene, often resulted in the highest CT values for the F gene amplification, indicating a low abundance of transcripts. Given its location in the genome, F should be a mid-abundance transcript. Again, data suggests transcript abundance may be non-linear that is strain dependent and thus F may not necessarily be mid-abundance (50). Increasing the temperature of the initial RNA reverse transcriptase hexamer binding resulted in in lower CT values, and better detection of the F gene that were more in line with the rest of the RSV genes. These results suggested that some form of RNA secondary structure could be influencing the ability of DNA primers to bind the RNA affecting reverse transcription efficiency. Higher temperatures are capable of relaxing secondary structures, allowing the polymerase better access to the template and to increase reverse transcription efficiency. As such the genomic and mRNA secondary structure of the virus became of interest.

Surprisingly, little attention has been focused on the effects of RSV RNA structure (genomic or mRNA) in regulation of either transcription or translation of the virus. Nucleoprotein binds tightly to the genomic and anti-genomic strands making secondary RNA structures in them difficult to form. Thus, here, we concentrate on secondary RNA structure in the mRNA of RSV F transcripts. Multiple mechanisms controlling RSV translation including ribosomal frameshifting and shunting have been proposed (1,9,10). Currently, RNA termination-reinitiation in the M2 mRNA in this overlap region is the most accepted mechanism (11). Work on the M2 gene showed a conserved and essential region that was highly structured. The M2 mRNA transcript is translated into the two proteins M2-1 and M2-2 in a not well understood coupled translation mechanism. Due to the placement of the conserved structure, it is likely that the secondary RNA structure is causing the coupled translation event. This structure indicates that there could be other such RNA secondary structures within other mRNA transcripts, the genome, or the anti-genome that influence the regulation of gene expression in RSV (10).

While the work done on M2 is promising, there are precious few other papers looking into other possible sites of secondary RNA structures such as within F that might be influencing gene regulation in RSV. In other RNA viruses, like Dengue and Influenza A, a number of host binding proteins regulate viral replication by enhancing, blocking, or helping with RNA movement around the cell (12–14). However, their genomes are different from mononegavirales and not as highly encapsidated by the nucleoproteins. Previous analyses of influenza A virus (IAV; another negative-sense RNA virus) have discovered numerous structural elements that are implicated in critical processes (15). In the IAV genomic RNA long-range base pairing interactions form a “panhandle” secondary structure that is essential for forming the promoter for replication (16). In IAV coding RNAs, motifs have been discovered that regulate alternative splicing and that are critical to viral fitness (17) but IAV is different in that it replicates in the genome and thus any structures in RSV’s transcripts would not likely be for splicing. Other motifs have apparent functions in the regulation of translation (18) or that may be able to affect genome packaging (19).

While it is suggested that RSV’s M (20) and known that RSV’s M2-1, L, and N (genomic RNA only) proteins (21,22) are all self-viral RNA binding proteins (either directly or indirectly through interaction with another protein), not much is known about RSV’s RNA structure and host protein (if any) binding to genomic or mRNA. RNA folding is achieved by intra-molecular interactions between the nucleotides and are often sites of specific interactions with proteins. A similar approach to the one used recently for analysis of ZIKA and HIV genomes using ScanFold-Fold (23) was used across a multitude of RSV viral strains (A and B) to identify key sites of structural conservation. These sites may identify key sites of interactions with host or viral proteins critical to viral replication that could be manipulated to prevent or limit RSV infection. Again, caution with this statement should be taken as the genome is bound by the nucleoproteins.

In this study, we found a conserved viral RNA structure in the mRNA of the F gene through experimental and computational methods. This structure, which also is indicated computationally to have a mirror in the genomic RNA, appears to be critical in the RSV lifecycle as relaxation prevented viral rescue. Further understanding of RNA structures’ roles in regulating viral replication or protein or gene expression may foster therapeutic targets for this significant human pathogen.

## Materials and Methods

### Viral Strains

RSV A (A2, 2001, 2006, 1997, 1998, Memphis-37) and RSV B (B1) strains were obtained from Beiresources (Mananas VA) and expanded 1 time in HEp2 cells (ATCC, Mananas, VA). Virus was purified according to (24) with the following modifications. NT buffer was used to make 50% PEG 8000. 10% of the PEG solution was added to the viral supernatant and incubated with agitation at 4°C for 1hr followed by centrifugation at 5000xg for 40min. The resulting pellet was re-suspended in DMEM with 3% sucrose.

### Plasmid Construction

- **PCR2.1 containing the F gene-** Constructed using a TA cloning (Invitrogen, Carlsbad, CA, K204001) system according to manufacturer’s specifications and a PCR amplified F gene containing fragment.
- **Partial F Relaxed**- The relaxed F gene was generated by site mutagenesis of PjetF (25) in order to relax the paired hairpin within the middle of the F gene but retain a similar codon optimization as the original F. Specifically, the sequence ATTAAGCAAGAAAAGGAAAAGAAGATTTCTTGGTTTT at base pair 385 of F (within PjetF) was mutated to CCTGAGTAAAAAGCGCAAGCGCCGGTTCCTCGGCTTCTTC thus retaining the amino acid sequence TLSKKRKRRFLGFL.
- **PjetF-relaxed**- Constructed from the RSV PjetF described in (25) and a synthesized fragment of F containing the relaxed sequence. The mutated F fragment was cloned into the parent PjetF via HiBuilder (NEB, Ipswich, MA) and was sequenced.
- **pACNR–rHRSV-relaxed**- Constructed using the newly constructed PjetF-relaxed and the pACNR–rHRSV described in (25) via Hibuilder (NEB, Ipswich, MA) and sequenced.

### Scanning window analysis and structural motif discovery

The negative-sense RSV genomic RNA (NCBI: JQ901447.1), and it’s positive-sense coding RNA counterpart were both analyzed using ScanFold; a pipeline which consists of a scanning window analysis which is subsequently analyzed to highlight secondary structures which are most likely to be functional (26). The scanning window analysis (ScanFold-Scan) was run using a 120 nt window size, 1 nt step size, and several folding metrics were recorded (described in detail in (26). Two key metrics are the minimum free energy (MFE) and its associated z-score. The MFE was predicted with RNAfold (version 2.4.3) as it corresponds to the RNA secondary structure which would yield the lowest free energy during its formation (according to the Turner energy parameters (26).The corresponding thermodynamic z-score reports how unusual the MFE is for its particular nucleotide composition: by comparing the native MFE (MFEnative) to one hundred randomized sequences with the same mononucleotide content (MFErandom) and normalizing by the standard deviation of all MFEs (σ), thus indicating how many standard deviations more stable (i.e. negative z-scores) or less stable (i.e. positive z-scores) the MFEnative is than expected. Subsequently, ScanFold-Fold (26) was used to identify base pairs which consistently contributed to the formation of unusually stable RNA secondary structures. This is accomplished by generating a z-score weighted consensus secondary structure from all base pairs predicted across all analysis windows.

### Comparative sequence and secondary structure alignment

A total of 1222 Human RSV genomes were downloaded from NCBI Nucleotide database following a search for whole genomes for “Human orthopneumovirus”. Genomes were aligned to the ScanFold analyzed genome (accession JQ901447.1) with default settings using the MAFFT web server (27,28). The list of ScanFold-Fold identified base pairs (with Zavg < -2 from JQ901447.1) were compared to the alignment to determine the extent of structural conservation for each base pair throughout the alignment.

### qRT--PCR of Viral Genes

Virus RNA was extracted by Viral RNA/DNA isolation kit (Invitrogen, Carlsbad, CA) according to the manufacturer’s instructions. Primers for each gene were created on Primer-Blast (NCBI, Bethesda MD) and synthesized (IDT, Iowa City, IA). The sequences are as follows, Highly structured region (Primer1) F: gcaaagctgcagcatatcaa R: cattgttggacattaactttttctg Unstructured region (Primer 2) F: agaagtcttagcatatgtggtac R: aacagatgttggacccttcc RNA diluted at 1/100 was amplified using Luna® Universal One-Step RT-qPCR Kit (NEB, Ipswich, MA) according to the manufacturer’s directions on a QuantStudio3 (Applied Biosystems, Foster City, CA). These primers bind around the B structure in Figure 1.

**Figure 1:**
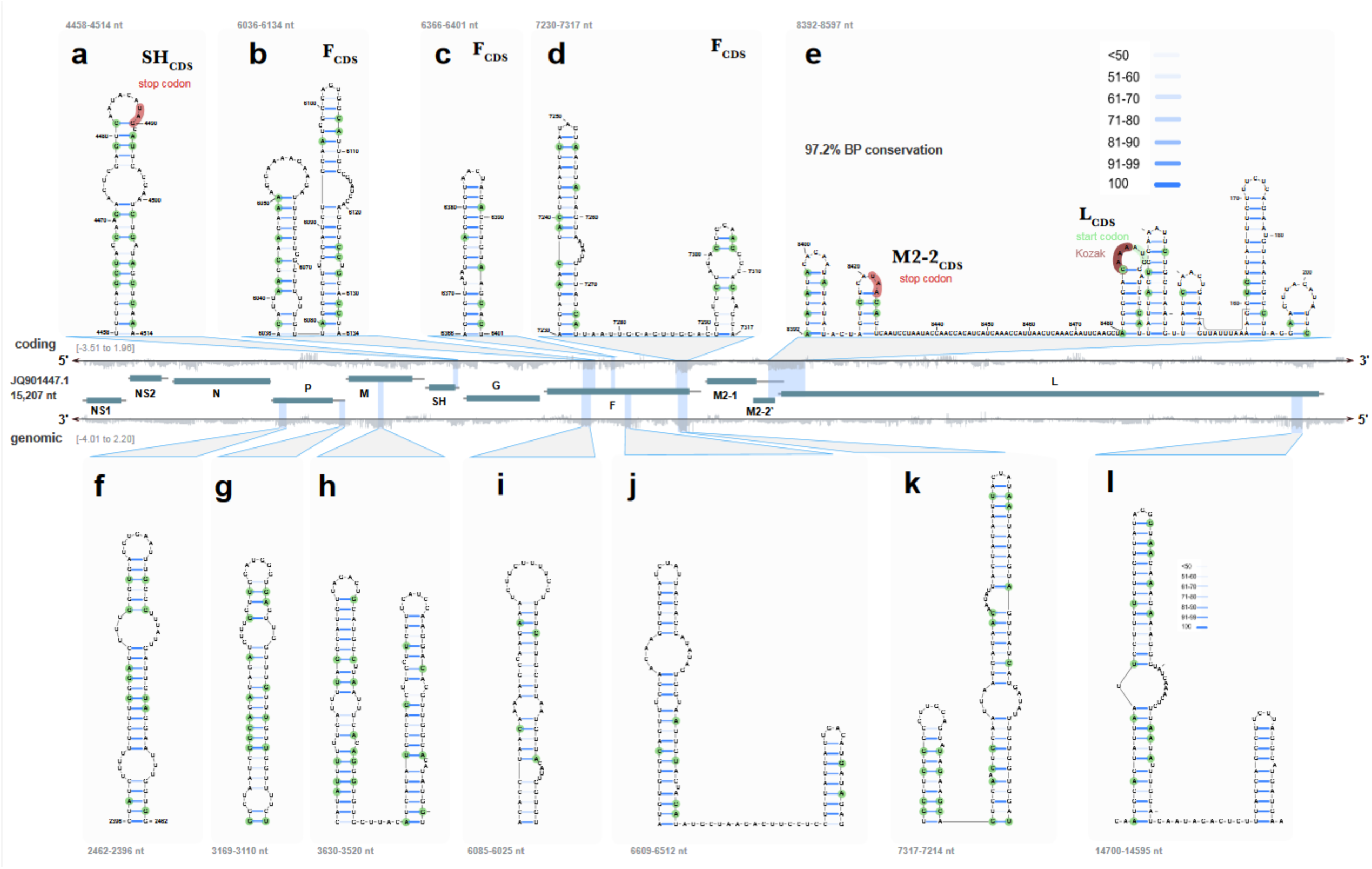
Scanning window/RNA structural motif discovery and analysis of the RSV genome and antigenome/mRNA transcripts. 1222 sequences of RSV were fed into ScanFold and areas of predicted (z-score >3) were plotted. Antigenome or mRNA transcripts are shown above the genome predicted structures.

### SHAPE Analysis of Predicted Structure

RNA was generated through InVitro Transcription according to (Invitrogen, Carlsbad, CA). 3µg of RNA was used for each SHAPE sample. After modification of the RNA was complete samples were reverse transcribed into cDNA using Superscript 3 according to manufactures specs (Invitrogen, Carlsbad, CA), with 1ul of a 1/10 of a 100uM stock reverse fluorophore tagged primer instead of the random hex/oligo dt. The sequencing ladder sample used a ddntpA mix in place of the dntp mix for use as a ladder. Protocol from here is as described in (29) re-suspending the final pellet in 10ul. Reverse primer sequence is as follows Set 1: cagccttgtttgtggatagtagag tagged with FAM. Samples were sent to be sequenced by capillary electrophoresis and then analyzed using QuSHAPE and RNAFold (30,31).

### Generation of SHAPE Figure

Done by mapping reactivity values onto the RNAFold structure generated from the SHAPE reactivity values or the ScanFold generated structure based off the models in (26) via RNA2Drawer (32).

### PCR of SHAPE Products

2ul of SHAPE produced cDNA and of cDNA produced from an unmodified FmRNA and cellular mRNA using reverse primers designed to target structured and unstructured regions were PCR’d according to manufacturer’s specifications (Takera, Mountain View, CA). PCR products were then run on an agarose gel. Primer sets are as follows, for cDNA made with the primer for the structured region Primer3 F: gtgttggatctgcaatcgcc R: gacttcctcctttattattgacatga with the unstructured region primers Primer4 F: agaagtcttagcatatgtggtac R: tttattggatgctgtacatttggt. Primer3 created cDNA was run at 60°C as was Primer4 as per required TM for primer sets for 20 cycles.

### Cell Transcription and Transfections

PCR products were generated using the PjetF and PjetF-relaxed as templates and the following primer set which contains a T7 promotor sequence Forward: aaataatacgactcactatagggccatggagttgccaatcctcaaaacaaatgc Reverse: tcagctactaaatgcaatattatttataccactcagttgatcc. PCR products wildtype and relaxed were then transfected into T7 expressing Baby Hamster Kidney Cells (33) (BHK: BSRT7) using the X2 transfection system according to manufacturer specifications (Mirus, Madison, WI). Transfections were incubated overnight, and cells were harvested. mRNA was extracted using RNeasy Plus according to manufacturer specifications (Qiagen, Hilden, Germany). Extracted RNA was treated with DNaseI to remove any residual contaminating DNA according to manufactures specifications (Thermo Fisher Scientific, Waltham, MA). Samples and standards were then amplified using the Luna 4x UDG system (NEB, Ipswich, MA) and the following primer set (Forward: tgcagtgcagttagcaaagg Probe: FAM/cagaattgc/ZEN/agttgctcatgca/ 3IABκFQ Reverse: gattgttggctgcttgtgtg) according to the manufacturer’s directions on a QuantStudio3 (Applied Biosystems, Foster City, CA). Copies per ul were calculated using Linear interpolation and the cts generated by the standards.

### In vitro transcriptions and RNA pulldowns

Plasmids containing either the RSV F in wildtype sequence or relaxed were used to generate RNA using biotin labeled UTP at 1/3 the amount of unlabeled UTP in a Biosearch Technologies (Middlesex UK) transcription kit. To minimize contributions from other structures or the UTRs, we only generated short transcripts (still predicted to fold correctly) around our predicted structure in region B of figure 1. RNA was purified by RNA purificiation kit (NEB, Boston MA). RNA was tranfected in Hep2 cells using RiboJuice (Millipore, Temecula CA) according to their instructions. Cells were then fixed with UV using a stratalinker and cells disrupted by sonication in the presence of protease inhibitor (HALT, Thermofisher, Waltham MA). Anti-bioin magnetic beads were reacted and RNA pulled down by magnet. Pull-downs were added to PAGE gel loading dye and boiled before loading on PAGE gels (Invitrogen, Carlsbad CA), transferred, and blotted using antibodies (Genetex, Irvine CA) to identified binding proteins.

### Imaging of PCR Transfections

Duplicate samples of those harvested for RNA were harvested and fixed in solution. Cells were stained for F using MPE8 (5) and an anti-human Alexa Fluor 555 secondary. Imagining was done by dotting a small amount of the stained cell solution on a slide and using a fluorescent scope.

### Construction and Rescue of RSV using Reverse Genetics

This was done according to prior published studies (25) with the following modifications: Initial transfections were done using the X2 system and were scaled up 2.6x and dropped on to slightly under-confluent BSRT7 cells in a 25cm2 flask. Both pACNR–rHRSV-relaxed and pACNR–rHRSV transfections were left on cells for 4hrs before cells were washed twice and new media was added to the transfected cells. Transfected cells were incubated for ∼4 days or until media color indicated it needed to be changed or cells became 100% confluent. Both virus and infected cells were harvested and moved to a larger flask after this time. Virus was sucrose purified after a final incubation in a 185cm2 flask and 200ul were taken for viral RNA extraction and to be plated onto new HEp-2 cells for staining.

### Intracellular Staining of wildtype and relaxed Virus

Viral infection was incubated for 24hrs before cells were washed and fixed in paraformaldehyde. Cells were washed in 1x Permwash twice and incubated in Permwash for 30 min. at 4°. Cells were stained for RSV using a human anti-RSV antibody (Beiresources, Mannasas VA) and an Alexa Fluro 555 secondary (Invitrogen), were re-fixed and imaged.

### Surface Staining of wildtype and relaxed Virus

Infected BSRT7 were harvested at the same time as viral harvest and were surface stained for F using the MPE8 (51) antibody and anti-human Alexa Fluor 555 secondary. Imagining was done by dotting a small amount of the stained cell solution on a slide and using a fluorescent scope.

### Copies per ul for wildtype and relaxed Virus

Viral RNA was extracted using the Nucleospin system according to manufactures specifications (Macherey-Nagel, Düren,). Extracted RNA samples and DNA standards were then amplified using One Step PrimeScript™ RT-PCR Kit (TAKARA BIO INC, Kusatsu, Shiga, Japan) and the following primer set ((Forward: atggacacgacacacaatga Probe: FAM/ccacaccac/ZEN/aagactgatgatcaca/ 3IABκFQ Reverse: gtggcctgttttcatcaag) according to the manufacturer’s directions on a QuantStudio3 (Applied Biosystems, Foster City, CA). Copies per ul were calculated using linear interpolation of the CTs generated onto a standard curve of a plasmid containing the NS1 gene.

### Statistics

Student T tests were run on the F gene expression of plasmids and viral infection for predicted structure and unstructured regions. Student T tests were run on the mRNA transfections between log copies per ul of wildtype vs relaxed mutants. Student T tests were run on the log copies per ul of viral concentration recovered from wildtype vs relaxed mutants. A p-value of less than 0.05 was considered significant.

## Results

### Scanning window analysis

The RSV genome was analyzed using the ScanFold pipeline with a window size of 120 nt and step size of 1 nt. This resulted in 15088 overlapping windows spanning both genomic and complementary (anti-genome/mRNA) RNAs (Figure 1). For each window several RNA folding metrics were predicted, which can be used to assess the sequence’s capacity to form an RNA secondary structure. Two metrics of particular interest are the predicted minimum free energy (MFE) of the window sequence and its associated z-score. The resulting MFE and z-score values for each window are plotted vs. the RSV genome in Figure 2. Across the coding sequence, native MFE values range from -36.7 to 0.0 kcal/mol, averaging -15.67 kcal/mol; across the genomic sequence native MFE values range from -40.4 to -3.4 kcal/mol, averaging -18.98 kcal/mol. Across the genome, thermodynamic z-scores ranged from -4.68 to 2.57 and -5.19 to 2.50 in the coding and genomic strands respectively. Despite having a more positive max and min z-score, the average thermodynamic z-score of the sequence was slightly more negative (-0.44) than the genomic sequence (-0.36) suggesting that overall, the coding sequence has more bias for being ordered (by evolution) to form stable secondary structures—indicating potential function. This was also observed in IAV (35) where regulatory elements were preferentially found in the coding RNA vs. the genomic RNA, which does not undergo complex regulation.

**Figure 2:**
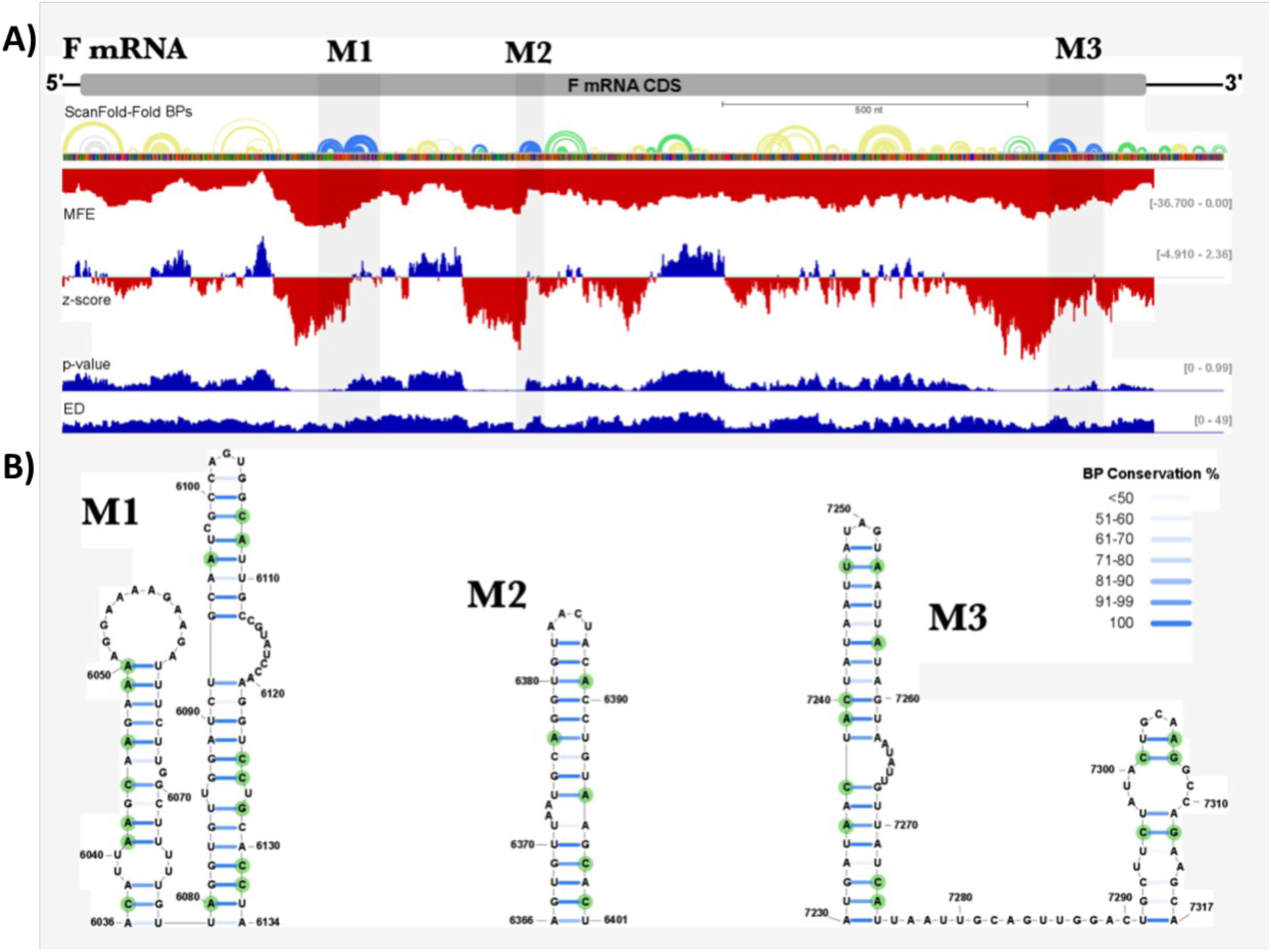
Scanning window/RNA structural motif discovery and analysis in the F mRNA region. **A)** ScanFold generated MFE and z-score values plotted against the RSV mRNA. **B)** Conserved structure predictions for the F mRNA similar to figure 1.

### RNA structural motif discovery

In order to determine the specific structures that could be giving rise to low z-score regions, the ScanFold-Fold algorithm (26) was employed. This algorithm systematically analyzes the MFE structure and z-score from each window and identifies base pairs which were consistently present in low z-score windows. In the strand, 29 RNA secondary structures were predicted to contain at least one base pair which consistently appeared in windows averaging z-scores less than -2 (Zavg< -2); with only 13 structures (greater than 4 bps in length) composed entirely of Zavg<-2 base pairs (Figure 1). These z-score generating structures comprise five structured regions which are found within four RSV mRNAs (here, structures are considered part of the same region if they are within 120 nts of each other). The first structured region consists of a single hairpin which spans the coding and 3’ untranslated region (UTR) of the SH mRNA. The next three structured regions occur throughout the coding region of the F protein mRNA. The fifth structured region is relatively large (205 nt long), spanning three different mRNA regions, and consisting of 7 hairpins composed entirely of Zavg<-2 bps and (ranging from 14 bps to 3 bps in length). This region is remarkable for its overlapping genetic roles; whereby it comprises the 3’UTR of M2-1, the stop codon for M2-2, and the 5’UTR and start codon for the L protein. This region also had the highest level of base pair conservation at 97.2% (as opposed to an overall average of 90.0% for all Zavg<-2 base pairs).

In the genomic strand of negative polarity, 20 RNA secondary structures were predicted to contain at least one bp with Zavg< -2. Of these, 14 were composed entirely of Zavg< -2 bps. comprising seven structured regions throughout the genome. The majority of these structured regions were located in different locations when comparing the genome locations and the coding sequence (Fig. 1 F, G, H, J, and L) with only two regions mapping to structured region locations in the coding strand (Fig. 1 I and K).

Since we initially were having different CT values detected when trying to amplify off the same cDNA strands for the RSV F mRNA, we decided to look more indepth whether conserved structures in this region might be impeding our amplifications of this transcript. As shown in Figure 2, the RSV F mRNA region (and mentioned above), had three conserved regions across all strains of RSVa.

### Template dependent amplification shift indicative of RNA Structure

Using the data obtained above, primers were designed to target the regions of 6273-6478 (Primer1) and 6528-6647 (Primer2) of the F transcript. The region targeted by Primer1 being predicted to be highly structured and Primer2 region being to be lightly structured or unstructured. The primers were used to generate cDNA from HEp-2 cells infected with RSV, uninfected Hep2 cells, and plasmid DNA from a PCR2 plasmid containing the F gene. The primer binding site had an effect on the degree of PCR amplification (Fig 3). Infected Hep2’s showed a difference in amplification with the Primer1 located in a region having higher CT’s than that of the Primer2 region. Higher ct’s correspond to a lower amplification/gene expression, while a lower ct correlates to a higher amplification/expression. Differences in the amplification of the two regions were significant as seen in the difference in relative fold increase over universal uninfected cells. Amplification on our RSV F plasmid showed no significant difference in the difference in relative fold increase over universal uninfected cells between the two regions. The plasmid DNA is not only double stranded and thus not structured, but was never subject to the inefficiencies in the reverse transcriptase that the viral cDNA was. Because of this, the plasmids should not have a significant difference between the two primers as there is no difference in the difficulty in amplification between the two sections that was predicted to occur in the viral RNA conversion to cDNA. This shows that in regions of predicted as structured there is a significant difference in amplification, not attributed to primer efficiency, giving more credence to the presence of a structured region in the F gene that may affect regulation of the gene.

**Figure 3.**
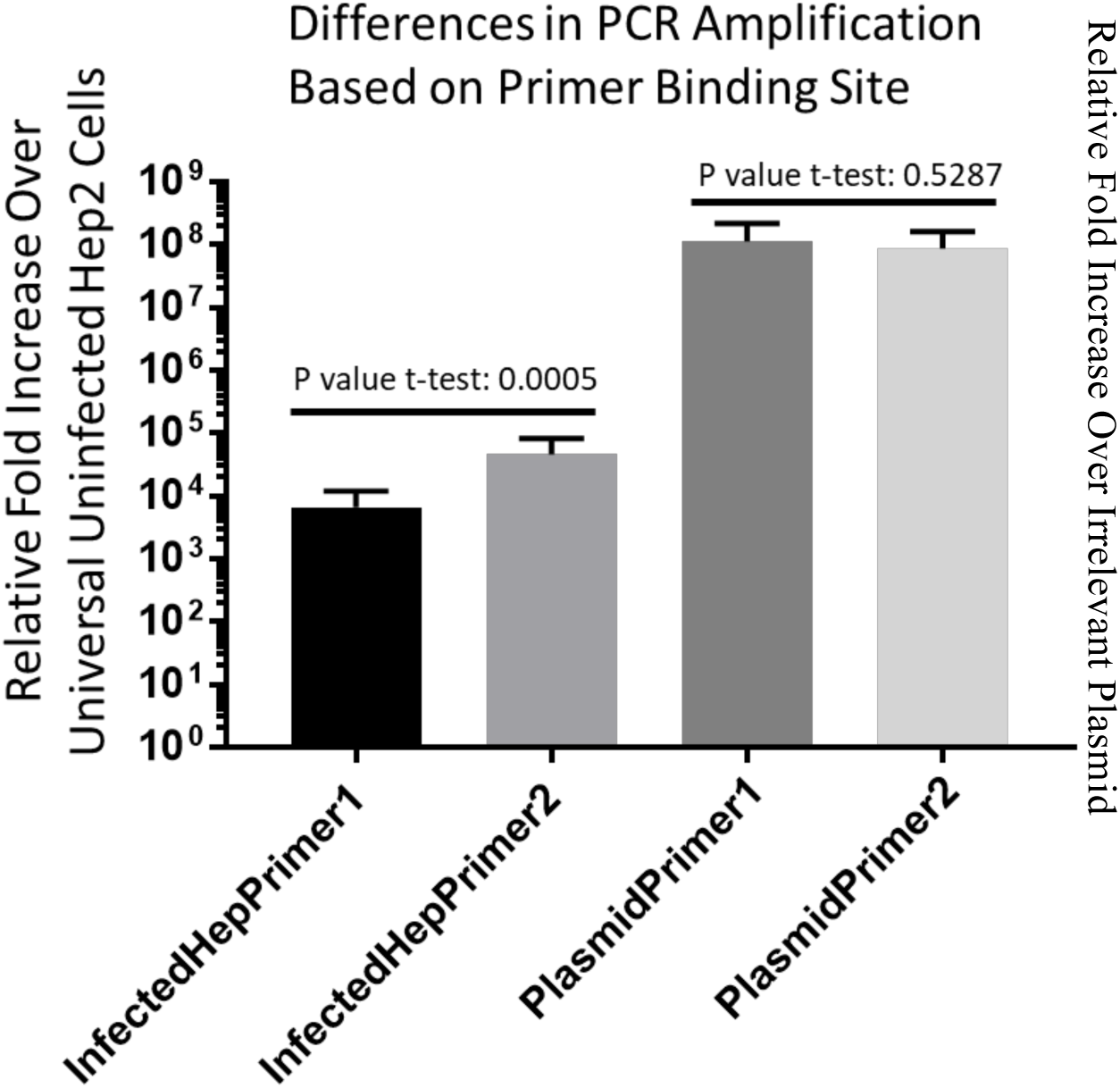
Primer binding site has an effect on the degree of PCR amplification. Primer1 corresponds to a predicted highly structured region and the Primer2 corresponds to a predicted low/unstructured region.you should described how were done the experiments and analyzed the results. How were quantified the PCR products? Is it Q-PCR or gel analysis?

### Experimental confirmation of computationally predicted RNA structure in F mRNA

#### Modified and Unmodified cDNA are differently amplified

PCR products created from SHAPE generated cDNA confirmed that the cDNA created was indeed F and that the amount of full-length transcripts that were produced off of the modified RNA in the structured region was lower than that of the unmodified RNA of the same region as seen in Figure 3 This difference was not seen in the modified and unmodified RNA generated from the unstructured region. Variable amplification of cDNA through the structured region showed that the modified RNA had the lowest number of transcripts that ran through the structured region and that the ladder cDNA which was the same concentration as the cDNA from the modified RNA had more full-length transcripts as seen by the darker band. This was not the case for the cDNA created from the non-structured region where all of the created cDNA has similar amplification. This further gives evidence to the RNA being structured at the Primer3 location as modification is intended to create truncated cDNA products where structure exists on the RNA, and non-structured regions are supposed to be able to produce full length products. Stable RNA structure is also known to cause truncation issues due to the enzyme falling off of the template.

#### Structure determination

Experimental determination of F mRNA secondary structure was determined through SHAPE probing using benzoyl cyanide modified RNA. The primer set used was designed to target the region of 6036-6134nt of the F transcript that the ScanFold results predicted a stable and very conserved structure. Normalized SHAPE reactivity values close to zero indicate a low reactivity and a high propensity to form base pairs whereas high reactivity values indicate highly modified bases that are likely to be unpaired because the benzoyl cyanide is able to access the nucleotides for modification. Areas of double stranded RNA inhibit modification and thus have lower reactivity. For our purposes reactivity cutoffs were defined as follows 0-.4, .4-.6, >.6 and were taken to be unreactive (base-paired), moderately reactive (intermediate levels of base pairing), and highly reactive (unpaired) respectively (36–40). SHAPE analysis was performed four times and the final normalized reactivity values generated from QuSHAPE of all sets were averaged and run in RNAFold to generate a predicted structure using the generated SHAPE restraints. Reactivity values were obtained for more than 90% of nucleotides in the target region sequence. Said values were then mapped onto the RNAFold and ScanFold generated structures in RNA2Drawer (32) to determine the placement of the normalized reactivity values. While the exact base pairing predicted by each model is not identical both produce two hairpins directly adjacent to one another in the predicted region. Additionally, within the second hairpin both models show identical tips and a large bulged out region that are highly modified (Figure 4). The four individual SHAPE data sets were also independently run on RNAFold and 75% of them generated the bulge and 100% had the same tip (data not shown). The regions that differ on the first hair pin contain a stretch of U’s that may “breath” and allow the real structure to alternate between a more linear and double stranded orientation and a bulged out of single stranded conformation. RNA structures in their biologically relevant environments are rarely static and often change conformation (41,42). Overall, the experimental data supports the existence of two hairpin structures in the region predicted by the computational model.

**Figure 4:**
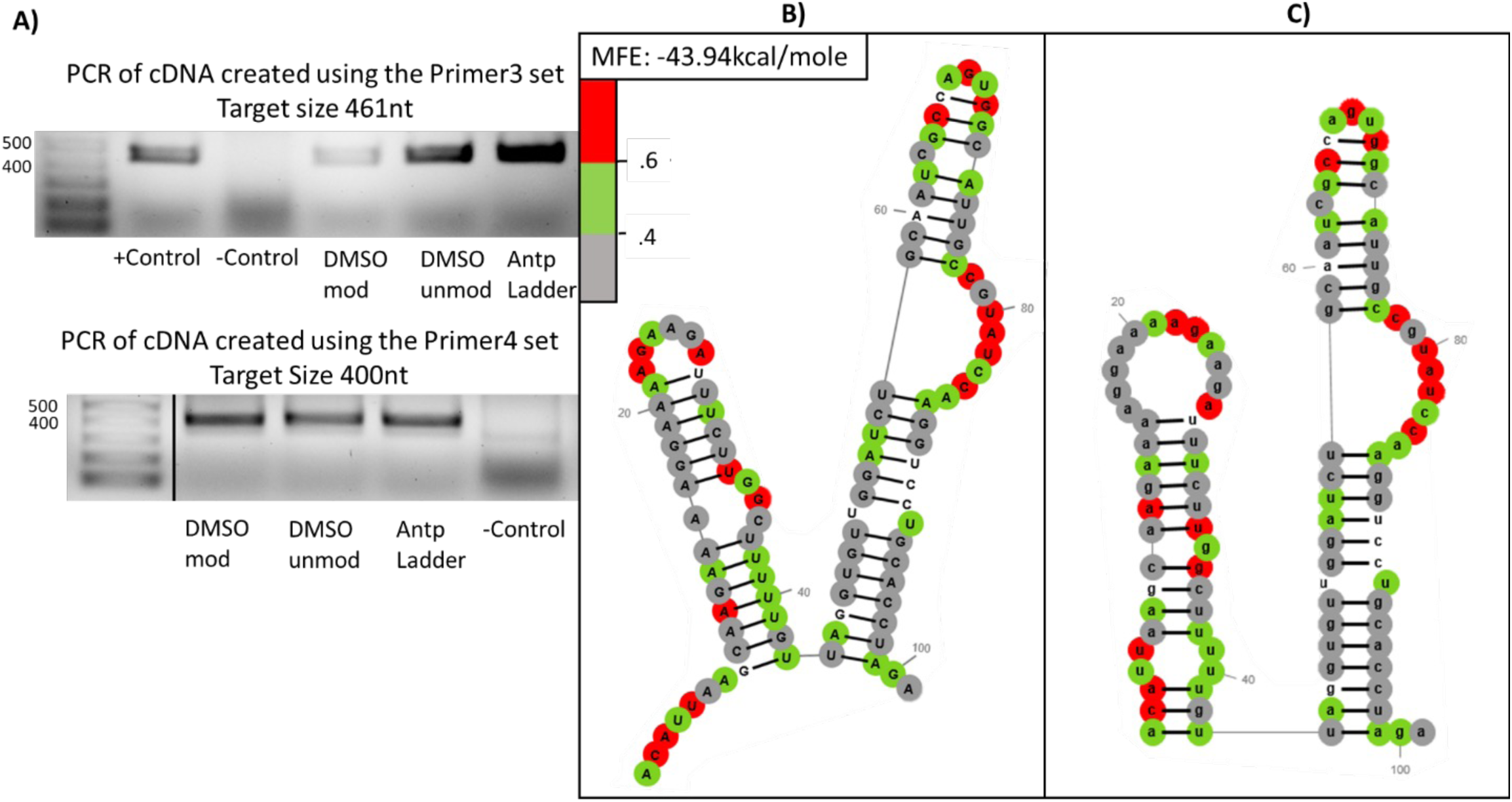
Experimental Determination of Secondary RNA Structure. **A)** PCRs of F off of SHAPE modified and unmodified cDNA using primers for structured and unstructured regions. Primer3 set is a region of predicted structure and Primer4 set is predicted to be unstructured. Modification is intended to create truncated cDNA products where structure exists on the RNA, and non-structured regions are supposed to be able to produce full length products. **B**) Normalized SHAPE reactivity values overlaid on the lowest MFE structure generated by RNAFold using the SHAPE reactivity restraints. 0-.4, .4-.6, >.6 were taken to be unreactive (base-paired), moderately reactive (intermediate levels of base pairing), and highly reactive (unpaired) respectively. **C)** Normalized SHAPE reactivity values overlaid on the computationally predicted structure shown in Figure 2 region M1.

### Production of F Protein Unaffected by RNA Sequence Change

First, we generated a relaxed version of the F gene on our plasmid to generate a change in mRNA structure at the same region as Fig 3 around the structure in region B of figure 1 using a shot-gun approach making a major change to the RSV structure located in this region. The amino acids remained the same and we tried to keep the codon usuage roughly the same.

To confirm that the modifications made to the RNA sequence of F and the subsequent relaxation of the secondary RNA structure in the mRNA did not affect the production and transcription of the F protein/mRNA we analyzed the levels of mRNA and protein produced from T7 BHK cells transfected with T7 promotor containing PCR products of either the wildtype F or the relaxed F (Fig 5A-C). The mRNA quantity was analyzed by qRT-PCR and indicated slightly higher levels of Relaxed mRNA were produced with slightly higher amounts of the relaxed version (Relaxed) being produced (Fig 6A). The copy number of produced mRNA correlated with the protein production of F as visualized by intracellular staining slightly higher in the relaxed version (Fig 6B).

**Figure 5:**
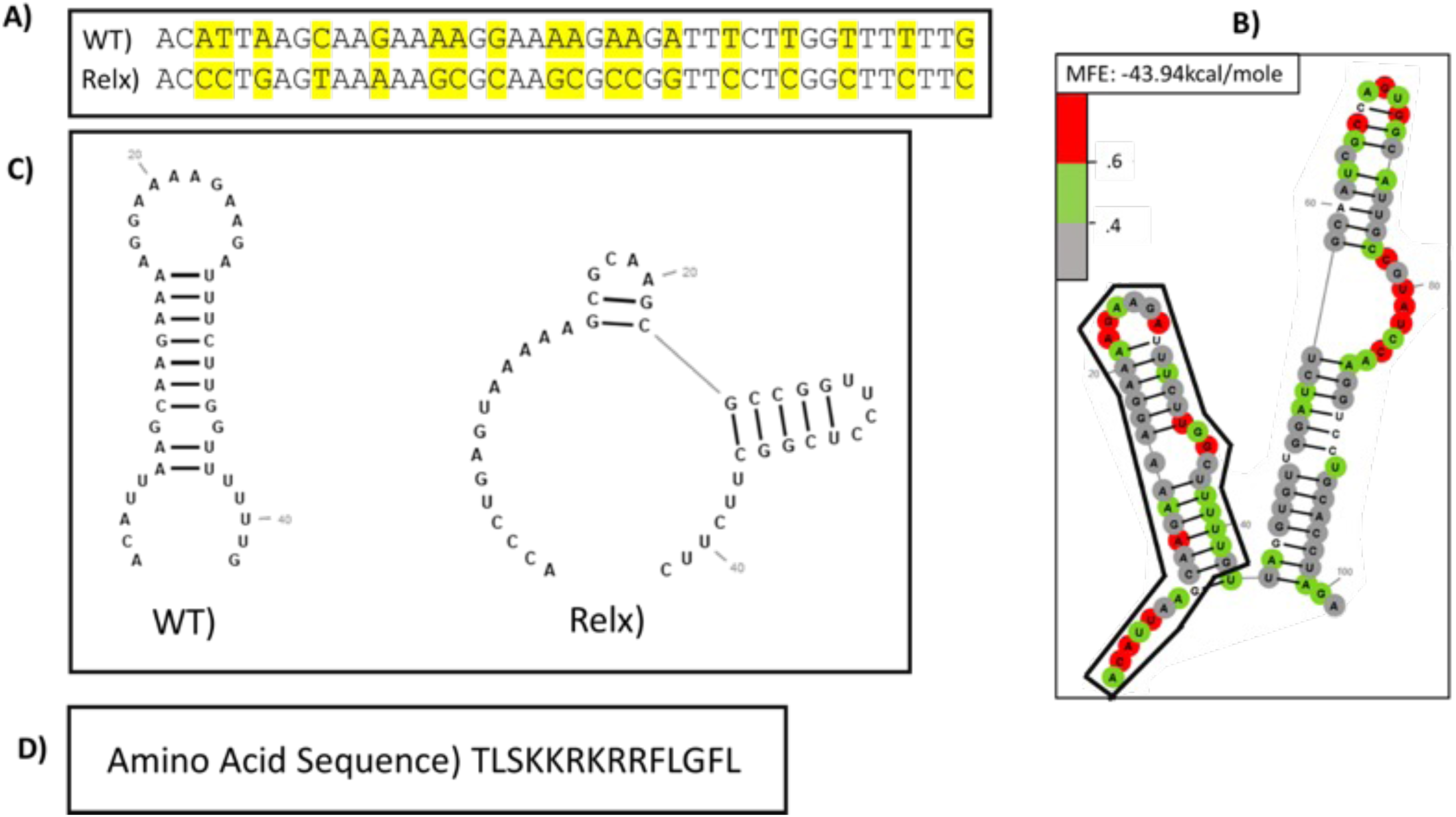
Design of relaxed mutation for F. We concentrated on the same M1 region form Figure 2 and determined the amino acid sequence. **A)** We then designed a relaxed mutant keeping the same amino acids and a similar codon usuage (Genscript table) **B)** but only focusing on ½ of the conserved structure (encircled in black). **C)** RNAfold prediction showed that the relaxed region should destroy the secondary RNA structure in RSV F mRNA. **D)** Interestingly, this region aligns with the highly conserved furin cleavage site in RSV F protein. WT: wildtype, Relx: relaxed mutant.

**Figure 6:**
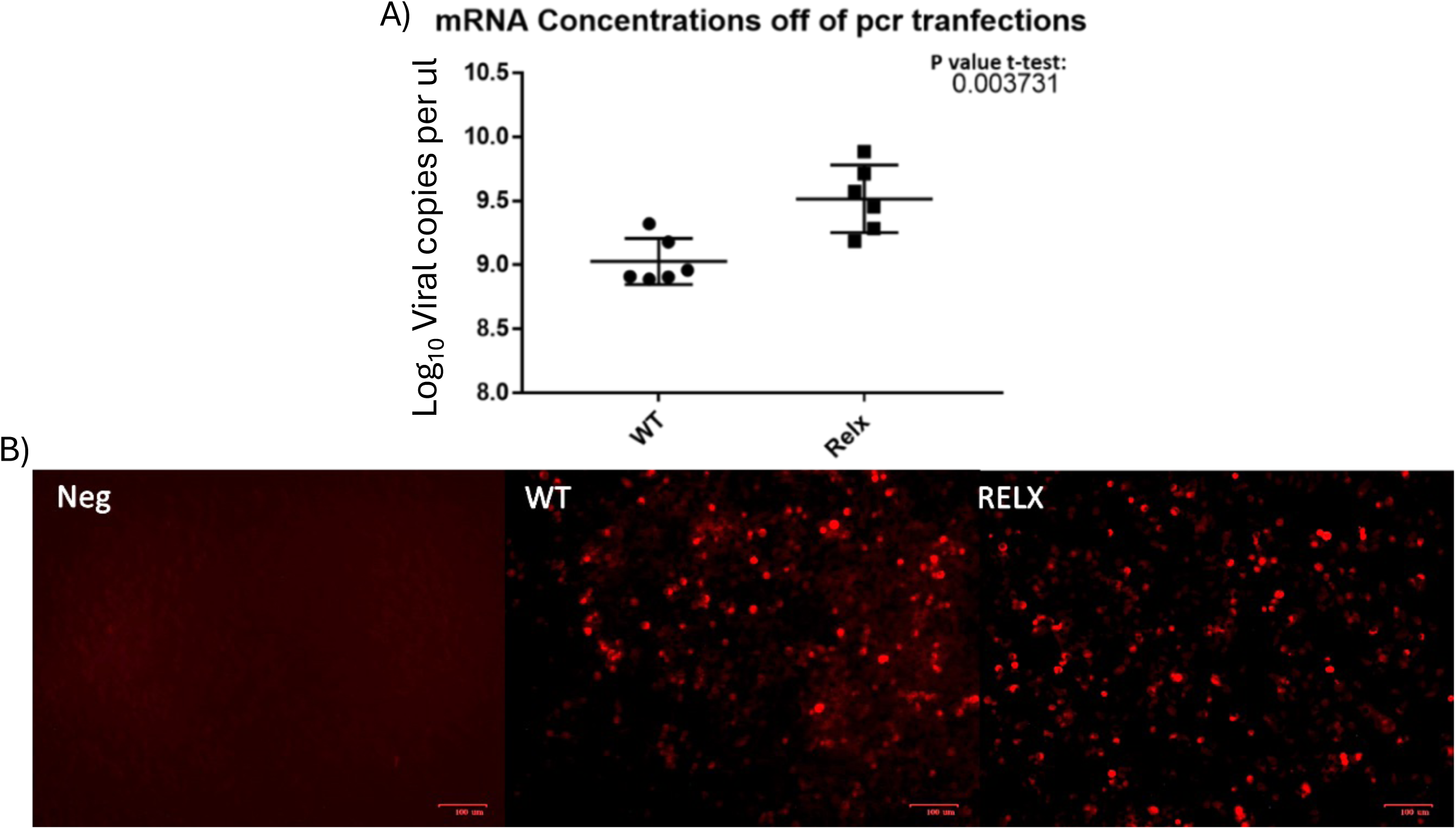
F Protein and mRNA of WT and Relx F. **A)** Calculated copies per μl on a log10 scale of mRNA harvested off of either PCR fragment wildtype or relaxed transfected T7BHK cells. **B)** Anti-RSV stained cells transfected with either wildtype or relaxed F PCR fragments. Red is detected virus.

### RNA Structure relaxation prevents viral rescue

To further test whether altering the secondary structure would have an effect on viral production given that the T7 polymerase system is very robust at generating mRNA off DNA with the correct promoter, we used the same wildtype, relaxed region, and a region with a compensatory mutation in our shuttle vector that we then moved into a plasmid containing the viral genome (Fig 7A). Initially produced viral titer is very low and often below detection. As such we not only passaged the virus containing supernatant but the initially transfected cells so that any infected cells had the optimal opportunity to produce virus and to allow infection to spread to neighboring cells. Redistribution of the cell monolayer by splitting cells into a new flask also allows the viral infection to better spread through the flask as infection clusters (cells that are infected and their immediate neighbors) are redistributed so that new infection centers can form, and the virus may spread to new uninfected cells. As such, the relaxed RSV virus was given all available opportunities to rescue both in time of incubation and repeated passaging in multiple flask sizes. While the wildtype virus consistently cased syncytia and spread, the relaxed failed to do so. After ∼ 6 days the wildtype infected cells blew out causing a lifting of the monolayers and large quantities of syncytia cells were seen floating in the media. In contrast the relaxed infected cell monolayer as intact and the cells appeared only slightly stressed as did the negative control cells likely due to the pH of the media (data not shown). Dectection of virus after transfer of viral supernatant on uninfected cells after transfection of the reverse genetics system, was done by intracellular antibody staining. We could see high levels of RSV within cells in our compensatory mutant and wildtype viruses but much lower levels of F staining in our relaxed F rescue (Fig 7B). We then took the cell supernantants to determine how much virus was being released between the wildtype and relaxed and found a significant difference in virus with relaxed failing to replicate and release (Fig 7C). These data were very interesting as we orgnially thought from Fig 6 that relaxed would have lead to higher viral production.

**Figure 7:**
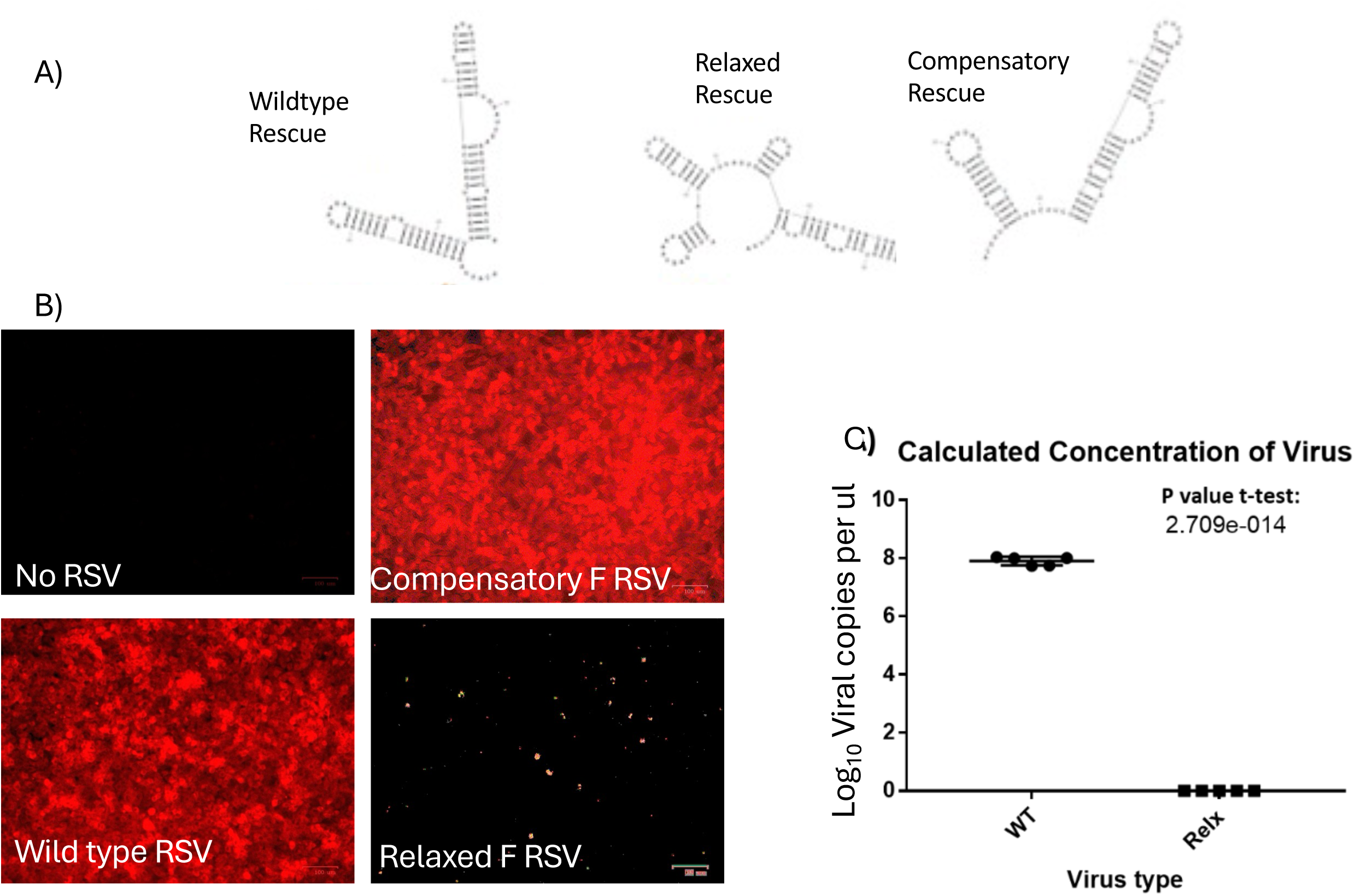
Relaxed RSV F mRNA impairs viral pathogenesis after viral rescue. We used our wildtype virus, our relaxed virus, but also generated a compensatory mutant in the same RSV region to restore the RNA structure to the same shape. **A)** RNAfold predicted structures are shown for comparison. **B)** We then used our reverse genetics system to rescue virus after transfection and transfer virus to new uninfected Hep2 cells. These were stained intracellularly with anti-RSV antibody and a fluorescent reporter. **C)** Viral supernantants were harvested and subjected to qRT-PCR to determine viral copy numbers. Neg: uninfected cells, other abbreviations similar as prior figures.

### Protein binding prediction identified a number of potential partner proteins interacting with this region of the F mRNA

We fed the structured region into RBPsuite and found a number of predicted proteins that could bind to this region of the RSV F mRNA (Figure 8A). We next generated RNA off our wildtype and relaxed F plasmids using in vitro transcription. We used biotin labeled UTP to allow us to capture the mRNA generated after we transfected the mRNA into Hep2 cells. After ruling out a number of predicted proteins (Pum2, MBL1 etc, not an exhaustive search), we probed on a western blot for EIF4B, we found a positive signal in the wildtype but not the relaxed site after magnetic pulldown on the RNA (Fig 8B). However, there could be other proteins binding that are subject to future studies. We only used a short version of the F transcript centered around the predicted folding and not be influenced by the parts of the mRNA (structured or not). Thus, EIF4B, which is expected to be on mRNA, would not have shown up in our experiment unless it was binding to our structure.

**Figure 8:**
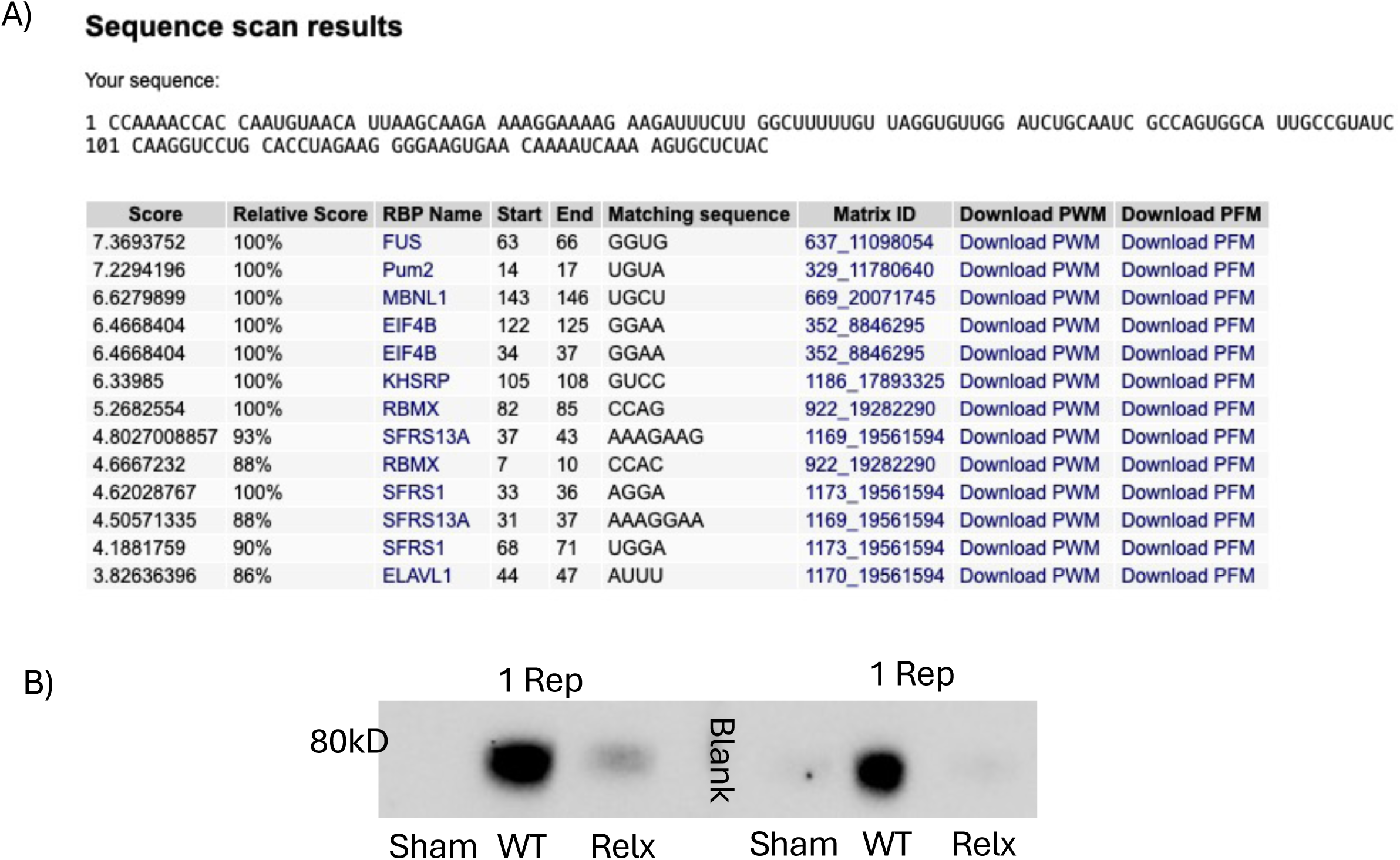
Predicted RNA binding proteins within the structured M1 region of RSV F mRNA. **A)** We used RBPsuite to feed in the sequence as outlined in the figure and generated a table of predicted proteins that could bind to these structured regions. **B)** RNA pulldowns after transfection of sham (unlabeled wildtype), wildtype, or relaxed F RNA suggest at least that EIF4B might bind to the region. Pulldowns were repeated 6 subsequent times and 2 representatives are shown in 1 blot

## Discussion

RNA secondary structure often serves an important regulatory purpose in mRNA transcripts, such as ensuring transcript stability and transport or influencing the timing or level of expression. Some secondary structures like those in the M2 mRNA of RSV are thought to allow access to alternative reading frames and allow for additional proteins to be produced through mechanisms such as ribosomal frameshifts. Other viruses use secondary structure to recruit RNA binding proteins which are crucial to their replicative life cycle (43,44).

ScanFold predicted the presence of conserved stable structures in nucleotides 6036-6134, 6366-6401, and 7230-7317 of the F transcript. This region corresponds to the coding region of the F gene, and is present in the F mRNA transcript. This region also corresponds to the fusogenic domain of F possibly explaining why this region may have more conservation than others. While the fusogenic domain region is important for the F protein and subsequently the conserved structure is on the RNA level. RSV has a high level of RNA sequence variation with the majority occurring in the G gene (45,46) but while sequence variability can affect protein viability, RNA secondary structure is not strictly dependent on the RNA sequence. Conservation in structure across viral strains often indicates a function within the viral lifecycle. The predicted structure information was used to design experimental tests.

qPCR data generated from primers designed to target predicted areas of structured and unstructured regions indicated the presence of a secondary RNA structure that inhibited the action of the reverse transcriptase resulting in a higher ct value than in areas of no structure. The higher ct of these regions was not found in the qPCR of plasmids containing the F gene indicating the difference was not due to primer inefficiency.

While the PCR amplification of the SHAPE generated RNA of the non-structured region was uniform regardless of whether the RNA was subjected to modification by benzoyl cyanide or not, the amplification of the SHAPE generated RNA from the structured region was not. cDNA off of the modified RNA amplified fewer full-length transcripts than that of the unmodified RNA or especially that of the ladder. The uneven distribution of full-length transcript amplification is further proof of the secondary structure existing within that region and that the SHAPE generated structure is indeed from F.

The structure found conserved in multiple RSV strains through ScanFold predictions was also found through experimental SHAPE probing giving high credibility to the presence and conservation of the structure. Highly conserved structures are often essential and thus can be targets for viral therapeutics. The F protein of RSV is an essential protein, as it allows for cell entry and is expressed on the viral envelope.

When this structured region of the RNA sequence was mutated to relax the secondary structure PCR transfection showed that the mutation did not alter the production of mRNA nor that of the F protein. This coupled with the failure to rescue the relaxed version of the reverse genetics RSV virus indicates that there is likely some regulatory function in the F mRNA lifecycle that the secondary structure serves as on its own the mutated sequence is able to produce F. While the mutation likely affects the mRNA, the computational analysis of both the coding and genomic sense of the F region indicates an almost twin structure located in the same region of F on the genomic strand. Mutation of the region to relax the mRNA structure would have also mutated the genomic structure. As such we can only say for sure that the relaxation of this structure in either/both the mRNA or/and the genomic region of F lead to the failure to rescue the relaxed virus.

The exact purpose of the secondary structure will be elucidated in further work but it is tempting to speculate what that purpose may be. Given that both the wildtype and relaxed mRNA are capable of producing the protein but the relaxed virus fails to rescue it is likely that the mutation causes a defect in a yet unknown regulatory mechanism that controls F expression, transcription, or trafficking within the context of a native infection. The RSV M2-1 protein may play a role as it is known to bind to nascent viral mRNAs not just at the polyA tail but also across the length of the transcript with a preference for A/U rich sites. Having bound the nascent mRNA, M2-1 can either leave the inclusion body and exit into the cytoplasm where it releases the mRNA for translation or go into a sub structure of the IB called an IBAG (52). The composition of viral RNAs and the duration of their stay are located in the IBAG, and as such it is a possible site of viral regulation. Perhaps the structure contributes in some way to a sequestration of the F mRNA in the IBAG. We are also unsure if there are more RNA binding proteins interacting with the structured regions in RSV F mRNA and do not quite know why EIF4B would be binding there since this translational helper usually binds IRESes or the beginning of transcripts. It is speculative but we would suggest that perhaps EIF4B might help the translational machinery stay on the transcript but given that relaxed T7 polymerase produced F had more expression this remains to be explored. Further work on exploring if EIF4B is needed or if other factors not yet identified but predicted are important to translation of F protein.

One caution we are aware of with our relaxed F structure work is the shift from adenine to guanine or cytosines. These subsititions could affect internal methylation of the mRNA and thus its stability. Further studies to slowly walk nucleotide point mutations are underway.

While we have examined potential RNA folding the RSV genome, nucleoproteins are known to tightly associate with the RNA and might prevent these RNA stuctures from forming. However, secondary RNA structures are known to occur in influenza virus, albeit it’s a segmented virus for which RSV is not, and nucleoproteins are tightly associated with that genome as well (53–54). However, nucleoprotein binding the the influenza genome is not uniform and a similar thing may occur in RSV (55). Thus, much more exploration will need to be focused on validating these structures in the RSV genome and their role if found. Secondary RNA structures in mRNA as described here would not be interfered by nucleoproteins and are well known to occur. This is the first paper that we are aware of that assigns highly conserved RNA structures in the RSV F transcript and a role in pathogenesis. Many more structures are left to be explored.

